# Synthetic biogenesis of chromoplasts from leaf chloroplasts

**DOI:** 10.1101/819177

**Authors:** Briardo Llorente, Salvador Torres-Montilla, Luca Morelli, Igor Florez-Sarasa, Miguel Ezquerro, Lucio D’andrea, Eszter Majer, Adrian Troncoso, Alisdair R. Fernie, José A. Daròs, Manuel Rodriguez-Concepcion

**Affiliations:** Centre for Research in Agricultural Genomics (CRAG) CSIC-IRTA-UAB-UB, Campus UAB Bellaterra, 08193 Barcelona, Spain; ARC Center of Excellence in Synthetic Biology, Department of Molecular Sciences, Macquarie University, Sydney NSW 2109, Australia; CSIRO Synthetic Biology Future Science Platform, Sydney NSW 2109, Australia; Instituto de Biología Molecular y Celular de Plantas (IBMCP), Consejo Superior de Investigaciones Científicas-Universitat Politècnica de València, 46022 Valencia, Spain; Sorbonne Universités, Université de Technologie de Compiègne, Génie Enzymatique et Cellulaire (GEC), UMR-CNRS 7025, CS 60319, 60203 Compiègne Cedex, France; Max-Planck-Institut für Molekulare Pflanzenphysiologie, 14476 Potsdam-Golm, Germany; Consejo Superior de Investigaciones Científicas (CSIC), Barcelona, Spain

**Author notes:** Corresponding authors: BL MRC.

## Abstract

Plastids, the defining organelles of plant cells, undergo physiological and morphological changes to fulfill distinct biological functions. In particular, the differentiation of chloroplasts into chromoplasts results in an enhanced storage capacity for carotenoids with industrial and nutritional value such as beta-carotene (pro-vitamin A). Here, we show that synthetically inducing a burst in the production of phytoene, the first committed intermediate of the carotenoid pathway, elicits an artificial chloroplast-to-chromoplast differentiation in leaves. Phytoene overproduction initially interferes with photosynthesis, acting as a metabolic threshold switch mechanism that weakens chloroplast identity. In a second stage, phytoene conversion into downstream carotenoids is required for the differentiation of chromoplasts. Our findings reveal that lowering the photosynthetic capacity of chloroplasts and increasing the production of carotenoids are not just the consequence but an absolute requirement for chromoplast differentiation, which additionally involves a concurrent reprogramming of nuclear gene expression and plastid morphology for improved carotenoid storage.

## Introduction

Plastids comprise a group of morphologically and functionally diverse plant organelles capable of differentiating from one plastid type to another in response to developmental and environmental stimuli (Jarvis and Lopez-Juez, 2013; Sadali et al., 2019). Such plastidial conversions are essential to sustain many fundamental biological processes and largely contribute to cell specialization in the different plant tissues. Among the different plastid types, chromoplasts are of great importance in nature and agriculture because of their capacity to accumulate high levels of carotenoids, plant pigments of isoprenoid nature that provide color in the yellow to red range (Egea et al., 2010; Rodriguez-Concepcion et al., 2018; Sun et al., 2018). Carotenoids such as beta-carotene (pro-vitamin A) are health-promoting nutrients that animals cannot synthesize but take up in their diets. They are also added-value compounds widely used in cosmetics, pharma, food, and feed industries as natural pigments and phytonutrients (Giuliano, 2017; Rodriguez-Concepcion et al., 2018).

Chromoplasts differentiate from preexisting plastids such as proplastids (i.e., undifferentiated plastids), leucoplasts (i.e., uncolored plastids in non-photosynthetic tissues), and chloroplasts (i.e., photosynthetic plastids). Chloroplasts transform into chromoplasts during the development of many flowers and fruits, but only a few plant species differentiate chromoplasts in leaves (Sun et al., 2018; Sadali et al., 2019). The yellow to red colors that some leaves acquire as they senesce (e.g., in the autumn or when they are exposed to continuous darkness) are due to chloroplast carotenoids becoming visible when the chlorophylls degrade. This senescence process, however, does not involve the transformation of chloroplasts into chromoplasts but into a completely different type of plastids named gerontoplasts (Jarvis and Lopez-Juez, 2013; Sadali et al., 2019).

The most prominent changes during chloroplast-to-chromoplast differentiation are the reorganization of the internal plastid structures, together with a concurrent loss of photosynthetic competence and overaccumulation of carotenoid pigments (Egea et al., 2010; Jarvis and Lopez-Juez, 2013; Lado et al., 2016; Llorente et al., 2017; Sun et al., 2018; Sadali et al., 2019). The remodeling of the internal plastid structures generates an increased metabolic sink capacity but it also promotes carotenoid biosynthesis. The control of chromoplast differentiation appears as a very promising strategy for improving the nutritional and health benefits of crops (Lado et al., 2016; Giuliano, 2017; Llorente et al., 2017; Sun et al., 2018; Wurtzel, 2019). However, very few inducers of chromoplast development have been identified to date. Orange (OR) chaperones are among the best characterized, but they only work in some tissues and the specific mechanism by which they promote chromoplast differentiation remains unclear (Sun et al., 2018). The experimental manipulation of chromoplast differentiation for fundamental studies and biotechnological applications therefore requests a much better understanding of the mechanisms regulating this process.

## Results

### The activity of the bacterial crtB enzyme induces the transformation of leaf chloroplasts into plastids of chromoplast features

The first committed step of the carotenoid pathway is the conversion of geranylgeranyl diphosphate (GGPP) to phytoene, catalyzed by phytoene synthase (referred to as PSY in plants and crtB in bacteria). We previously found that the virus-mediated expression of a bacterial *crtB* gene in tobacco (*Nicotiana tabacum* and *N. benthamiana*), tomato (*Solanum lycopersicum*), *Arabidopsis thaliana* and many other plants caused leaf yellowing (Majer et al., 2017). This phenotype was deduced to result from a combination of virus-induced chlorosis and increased accumulation of colored carotenoids downstream of phytoene. To further study its possible cause, we analyzed plastid ultrastructure by transmission electron microscopy (TEM). Analyses of *N. tabacum* leaves treated with crtB-producing viral vectors showed that yellow sectors contained plastids with a distinctive ultrastructure that were absent in empty vector controls (Fig. 1A). These plastids were devoid of organized photosynthetic thylakoid membranes and grana but contained tightly packed membrane stacks and a proliferation of small electron-dense (i.e., lipid-containing) plastoglobules typically observed in chromoplasts (Fraser et al., 2007; Egea et al., 2010; Lado et al., 2016). Very similar structures were observed when the *crtB* gene was virally expressed in *Arabidopsis* leaves (Fig. 1B) or transiently expressed in agroinfiltrated *N. benthamiana* leaves (Fig. 1C). TEM examination of dark-incubated senescent *N. benthamiana* leaves confirmed that the plastids found in crtB-producing cells were completely different from gerontoplasts (Fig. 1D). Low expression of the senescence marker gene *SAG12* (Gan and Amasino, 1995) in *crtB*-expressing *N. tabacum* leaves further argues against these plastids being formed by a senescence process (Fig. S1).

**Figure 1.**
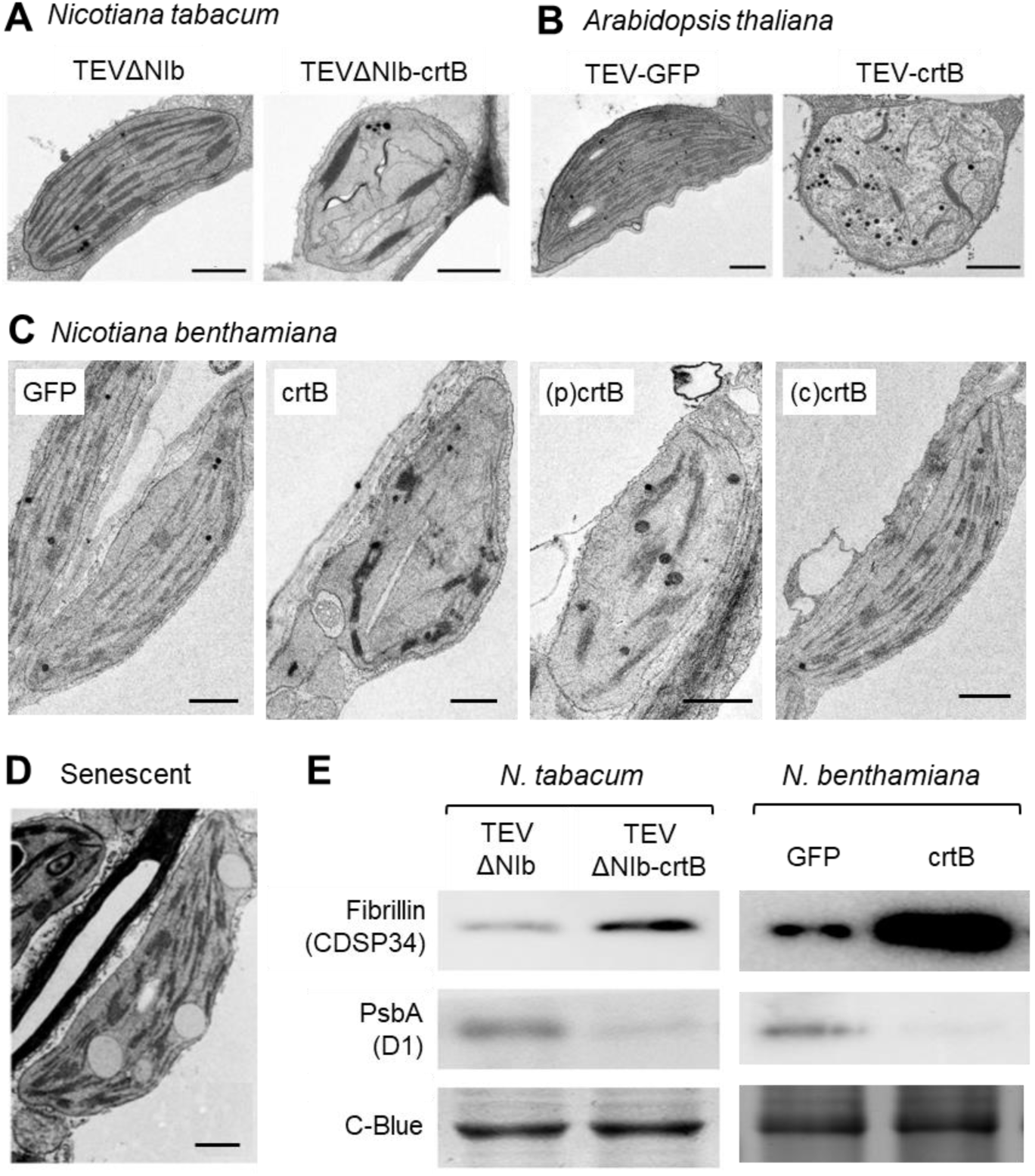
Chromoplast-like plastids develop from chloroplasts in leaves producing crtB or a plastid-targeted version of the enzyme. TEM images of representative plastids from the indicated species and treatments are shown. Bars, 1 µm. A, Plastids from *N. tabacum* leaves collected 10 days after inoculation with TEVΔNIb (empty vector) or TEVΔNIb-crtB. B, Plastids from *A. thaliana* (L*er*) leaves collected 15 days after inoculation with TEV (empty vector) or TEV-crtB. C, Plastids from *N. benthamiana* leaves collected 5 days after agroinfiltration of the indicated constructs. D, Gerontoplast from a *N. benthamiana* leaf harvested from the plant and kept in the dark for 10 days (senescent). E, Immunoblot analysis of plastidial proteins in leaves treated as described in panels A and C. Coomassie Blue (C-Blue) staining is shown as a loading control.

To further substantiate the identity of the chromoplast-like plastids that developed in crtB-producing leaves, we analyzed chloroplast and chromoplast marker proteins by immunoblot analysis (Fig. 1E). Virus- or *Agrobacterium tumefaciens*-mediated expression of *crtB* in *N. tabacum* or *N. benthamiana* leaves, respectively, resulted in increased levels of fibrillin, a protein associated to chromoplast development (Deruere et al., 1994; Singh and McNellis, 2011). In contrast, the levels of D1 (also known as PsbA), a core component of photosystem II (PSII) that is highly downregulated during chloroplast-to-chromoplast differentiation (Kahlau and Bock, 2008; Barsan et al., 2012), decreased in crtB-producing leaves (Fig. 1D). These results together suggest that expressing the bacterial *crtB* gene in leaf cells is sufficient to differentiate chloroplasts into chromoplast-like plastids.

We next used *Arabidopsis* double mutants defective in OR chaperones (AtOR and AtOR-LIKE) to test whether the differentiation process triggered by crtB involved pathways depending on these well-characterized promoters of chromoplast development (Zhou et al., 2015; Sun et al., 2018). Similarly, double mutants lacking cytosolic and plastidial carotenoid cleavage dioxygenases (CCD1 and CCD4, respectively) were used to investigate the possible contribution of signaling molecules derived from enzymatic degradation of carotenoids in the differentiation mechanism (Hou et al., 2016; Schaub et al., 2018; Wang et al., 2019). Virus-mediated expression of *crtB* in these mutants showed the characteristic yellow phenotype and carotenoid overaccumulation observed in the wild-type (Fig. S2), suggesting that signals derived from OR activity or from the enzymatic cleavage of phytoene or downstream carotenoids are not required for the process.

### The crtB enzyme only triggers chloroplast-to-chromoplast differentiation when localized in plastids

Consistent with the lack of a predictable plastid-targeting signal in crtB, a C-terminal fusion to the green fluorescent protein (crtB-GFP) mainly localized to the cytosol in *N. benthamiana* leaf cells (Majer et al., 2017). However, some fluorescence was also detected in chloroplasts. By optimizing the transient expression conditions we confirmed that the crtB-GFP protein was present in both cytosol and chloroplasts of agroinfiltrated cells (Fig. S3A). When GFP was fused to the N-terminus of crtB, the resulting protein (GFP-crtB) was completely excluded from chloroplasts (Fig. S3A). It is therefore likely that the bacterial crtB enzyme harbors a cryptic plastid-targeting signal in its N-terminus that becomes blocked and, hence, inactivated in the GFP-crtB protein. To unambiguously target crtB to the chloroplast, we next added the plastid-targeting sequence of the *Arabidopsis* enzyme HDS (Gas et al., 2009) to the crtB-GFP reporter. As expected, the resulting (p)crtB-GFP protein was only found in chloroplasts (Fig. S3A). Agroinfiltrated leaf tissues expressing either crtB or (p)crtB developed the characteristic yellow phenotype, whereas tissues expressing the cytosolic GFP-crtB version – renamed as (c)crtB – remained green as the controls expressing GFP (Fig. 2A). Analysis of carotenoid contents showed identical profiles for leaf sections agroinfiltrated with GFP and (c)crtB and confirmed that, similarly to crtB, (p)crtB triggered carotenoid overaccumulation (Fig. 2B). Unlike that observed with viral vectors in *N. tabacum* (Majer et al., 2017), however, agroinfiltration of *N. benthamiana* leaves with crtB or (p)crtB constructs did not reduce chlorophyll levels compared to leaf tissues agroinfiltrated with GFP or (c)crtB (Fig. 2B). Despite unchanged chlorophyll contents, estimation of photosynthesis-related parameters such as effective quantum yield of PSII (ϕPSII) and non-photochemical quenching (NPQ) showed that both crtB and (p)crtB, but not (c)crtB or GFP, had a dramatic impact on chloroplast function (Fig. 2C). A plastid-targeted version of GFP did not cause any yellowing or ϕPSII defect (Fig. S3B), confirming that disturbance of chloroplast photosynthesis is not caused by the accumulation of a foreign protein in chloroplasts but specifically by crtB. TEM analyses confirmed that (p)crtB induced the differentiation of chromoplast-like plastids very similar to those found in leaf tissues expressing the untargeted crtB enzyme, whereas only chloroplasts were present in leaves producing either (c)crtB or GFP (Fig. 1C). These results confirm that crtB elicits a synthetic (i.e., non-natural) differentiation of chromoplasts only when localized in plastids, where carotenoids are made.

**Figure 2.**
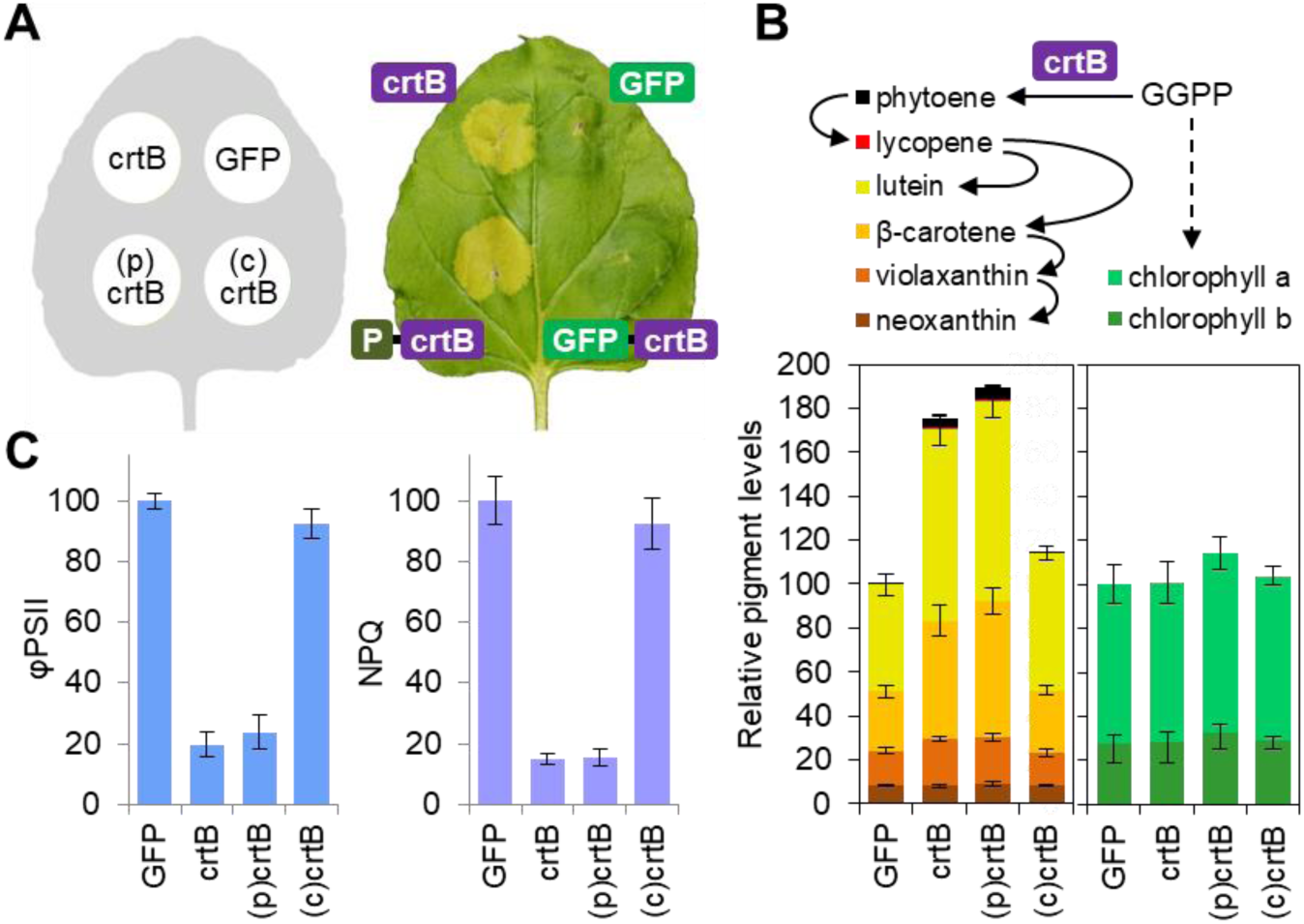
Bacterial crtB needs to enter the chloroplast to induce chromoplast differentiation. A, Phenotypes of *N. benthamiana* leaf sections 4 days after agroinfiltration with the indicated constructs. B, Photosynthetic pigment levels in leaf sections like those shown in A. C, Effective quantum yield (ϕPSII) and non-photochemical quenching (NPQ) in leaf sections like those shown in A. Plots show the mean and standard deviation of n=3 independent samples. Values are represented relative to those in GFP controls

### Synthetic chromoplast biogenesis induces profound changes in nuclear gene expression and primary cell metabolism

The vast majority of plastidial proteins are encoded by nuclear genes (Jarvis and Lopez-Juez, 2013). We therefore reasoned that the drastic remodeling of plastidial ultrastructure associated with crtB-triggered chromoplast differentiation would require changes in nuclear gene expression. RNA-seq analyses of *N. benthamiana* leaf samples at 96 hours post-infiltration (hpi) showed that about 5.000 genes were differentially expressed in yellow (p)crtB sections compared to green GFP controls (Table S1). Such a massive reprogramming of gene expression included the upregulation of 3.183 genes and the downregulation of 1,803 genes in chromoplast-containing samples. Gene Ontology (GO) term enrichment analyses (Table S2) showed overrepresentation of genes involved in protein folding and binding to RNA and ribosomes among those induced by (p)crtB (Fig. 3). Enrichment of genes with roles in transmembrane transport, cell signaling (protein phosphorylation, calcium binding), and nuclear gene expression (transcription factors) was observed among those repressed when chromoplast biogenesis was induced (Fig. 3). This profile was strikingly similar to that of ripening tomato fruits (where chromoplasts naturally differentiate from chloroplasts) but very different from that of senescent *Arabidopsis* leaves (Fig. 3). In the case of crtB-triggered leaf chromoplastogenesis, however, only minor changes were observed in the expression of genes with roles in carotenoid biosynthesis, degradation, or storage (Fig. S4 and Table S3).

**Figure 3.**
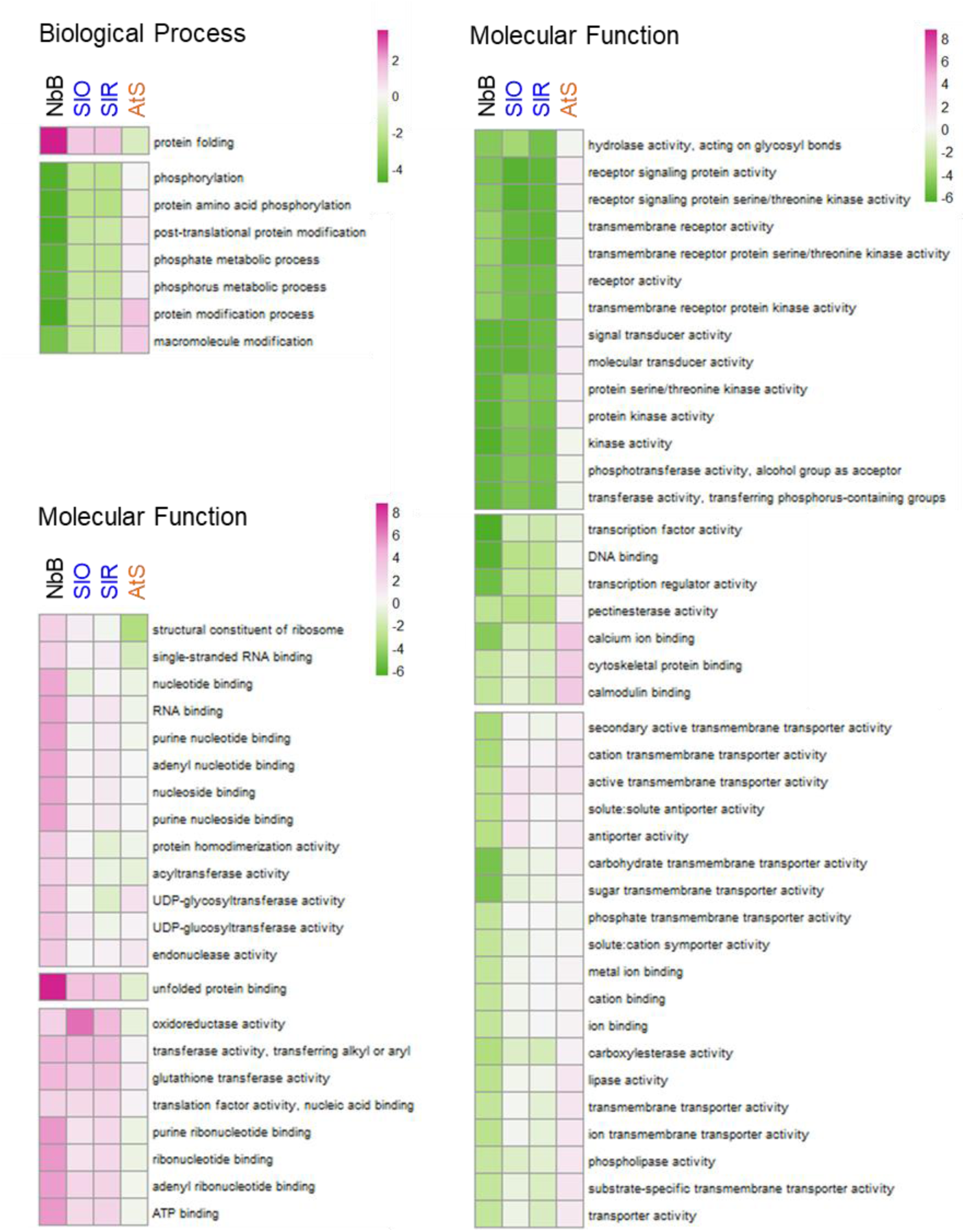
Chromoplast differentiation triggered by crtB in *N. benthamiana* leaves resembles that associated to ripening in tomato fruit. Heatmap represents GO term enrichment values (Z-scores) calculated from log2FC values in *N. benthamiana* leaves 4 days after agroinfiltration with (p)crtB compared to GFP (lane NbB). Data publicly available from tomato fruit (light ripe vs. mature green, SlO, and red ripe vs. mature green, SlR) and *Arabidopsis* leaves (senescent −30D- vs. controls −16D-, AtS) are also shown. Only GO terms with a significant enrichment in the *N. benthamiana* experiment are shown.

Energy and carbon required for carotenoid biosynthesis rely on photosynthesis (i.e., the Calvin-Benson cycle) in chloroplasts. Given that leaf chromoplast differentiation was associated with impairment of photosynthesis (Fig. 2C), we asked whether primary cell metabolism might also be reprogrammed. Of 52 metabolites detected by GC-TOF-MS analysis (Table S4), 13 displayed statistically significant changes in (p)crtB leaf sections compared to GFP controls (Fig. S5 and Table S5). We observed reductions in the levels of ascorbate and hexoses (glucose and fructose, the main soluble carbohydrate stores and respiration substrates). This, together with increments in tricarboxylic acid (TCA) cycle intermediates (citrate, 2-oxoglutarate and malate) and amino acids (valine, isoleucine, aspartate, and glutamate), suggested that sugars were used to produce ATP through the TCA cycle to sustain amino acid and carotenoid biosynthesis (and likely other cellular functions). Indeed, respiration rate – determined as total oxygen consumption in the dark – was higher in chromoplast-containing leaf tissues (Fig. S5). While an increased respiration is also associated to the onset of carotenoid overproduction in tomato and other climacteric fruits, the metabolic changes that we observed in (p)crtB-producing *N. benthamiana* leaves are often opposite to those occurring during chromoplast differentiation in tomato (Carrari and Fernie, 2006; Fraser et al., 2007). In particular, hexoses and ascorbate do not decrease but increase, whereas TCA cycle intermediates do not increase but decrease when tomato fruit ripens. We speculate that this might be because leaf metabolism is devoted to produce and export photoassimilates, whereas tomatoes are sink organs that have been selected to accumulate sugars and acids as positive taste attributes. In any case, our data together show that the activity of crtB in leaf chloroplasts is sufficient to trigger a deep reprogramming of nuclear gene expression and whole cell metabolism associated with the differentiation of chromoplasts, a plastid type that is not naturally found in tobacco or *Arabidopsis* leaves.

### Enhanced supply of phytoene in chloroplasts can interfere with photosynthesis

To investigate the dynamics of the crtB-dependent chromoplast differentiation process, we next followed the time course of *crtB* expression, phytoene production, and downstream carotenoid accumulation after agroinfiltration of *N. benthamiana* leaves with the (p)crtB construct. Transcripts encoding (p)crtB were reliably detected at 24 hpi and peaked at 48 hpi (Fig. 4A), whereas phytoene started to accumulate between 24 and 36 hpi and suddenly increased at 48 hpi (Fig. 4B). Downstream carotenoids began to increase at 48 hpi (Fig. 4B). To follow chloroplast membrane remodeling dynamics, we also monitored ϕPSII, NPQ, and D1 protein levels as estimators of photosynthesis, photoprotection, and photodamage. Both ϕPSII (Fig. 4C) and NPQ (Fig. S6) remained unchanged up to 36 hpi, and then decreased as the levels of both phytoene and downstream carotenoids increased (Fig. 4). The levels of D1 started to decrease later, between 48 and 60 hpi (Fig. 4D), likely as a result of photodamage. A higher temporal resolution analysis of both ϕPSII and carotenoid levels between 25 and 40 hpi showed that phytoene levels increased before ϕPSII decreased, whereas downstream carotenoids took a bit longer to accumulate (Fig. 4E). Taken together these results suggest that the crtB-mediated production of phytoene causes a disruption of the chloroplast photosynthetic functionality before carotenoids start to overaccumulate.

**Figure 4.**
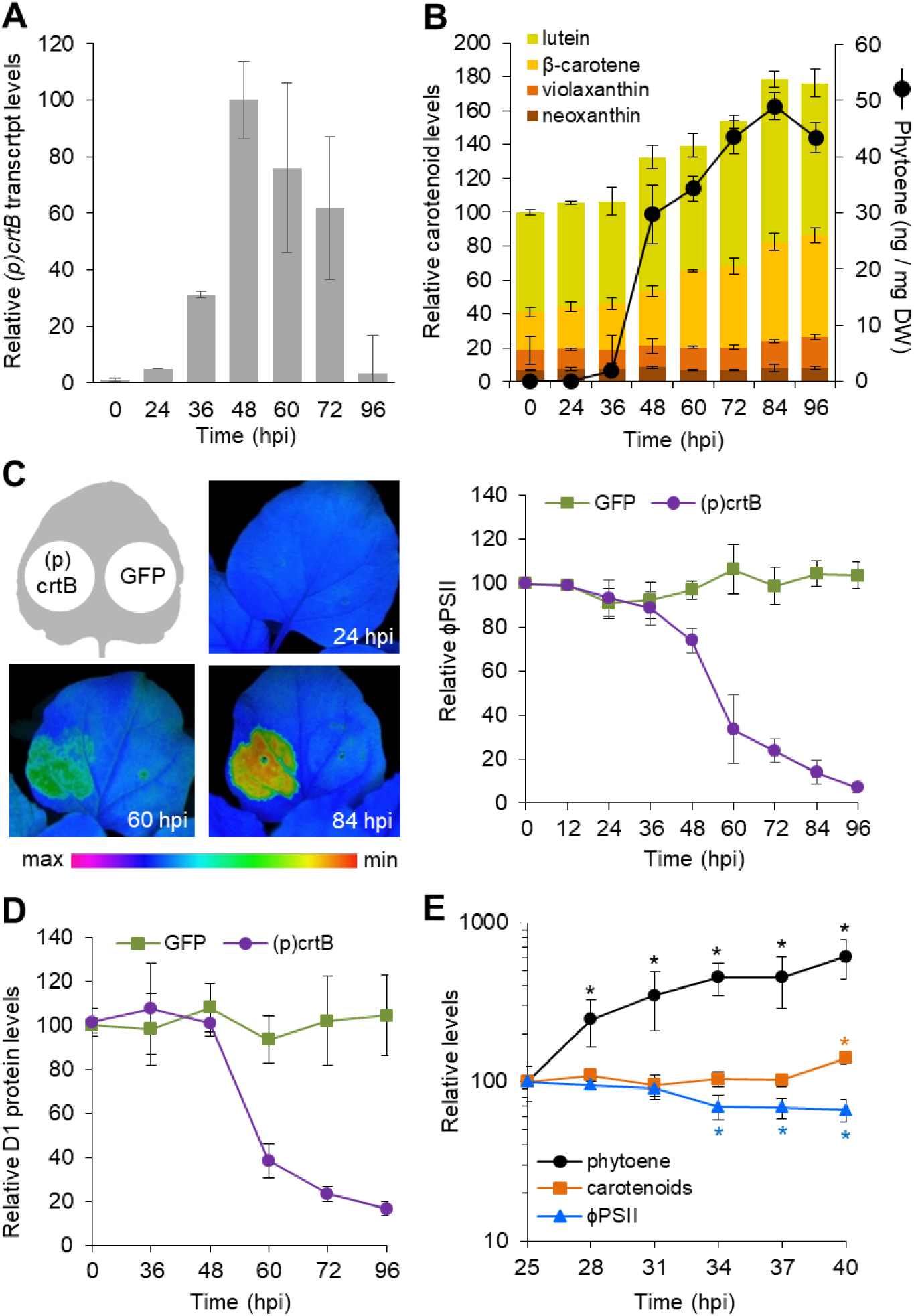
Time-course of chloroplast-to-chromoplast differentiation in leaves. *N. benthamiana* leaves were agroinfiltrated with the indicated constructs and samples were collected at the indicated time points (hours after agroinfiltration). A, Levels of (p)crtB-encoding transcripts relative to the maximum in (p)crtB samples. B, Absolute phytoene levels and relative downstream carotenoid contents in (p)crtB samples. C, Representative chlorophyll fluorescence images and ϕPSII values. D, PsbA (D1) protein contents. E, Levels of phytoene, downstream carotenoids, and ϕPSII in (p)crtB samples relative to those at 25 hpi. Asterisks mark significant changes relative to the 25 hpi values (*t* test, *P* < 0.05). Note the logarithmic scale. In all the plots, values correspond to mean and standard deviation values of n=3 independent samples.

To next confirm whether impairment of chloroplast functionality was due to phytoene overaccumulation, we used norflurazon (NF) to prevent phytoene conversion into downstream carotenoids (Ortiz-Alcaide et al., 2019). *N. benthamiana* leaves were agroinfiltrated with GFP and (p)crtB constructs and 24 h later some leaves were also infiltrated with NF (Fig. 5). The higher accumulation of phytoene in NF-treated (p)crtB relative to GFP samples correlated with a stronger reduction in ϕPSII (Fig. 5). By contrast, no changes in ϕPSII were detected in leaves transiently expressing the *Arabidopsis PAR1* gene, encoding a transcription cofactor that promotes total carotenoid biosynthesis but not phytoene accumulation in photosynthetic tissues (Roig-Villanova et al., 2007) (Fig. 5). We therefore conclude that it is the initial accumulation of phytoene but not of downstream carotenoids what causes a concentration-dependent disruption of the photosynthetic identity of the chloroplast. The observation that (p)crtB leaf sections treated with NF do not show the chromoplast-associated yellow phenotype and massive ϕPSII drop detected in the absence of inhibitor (Fig. 5) further suggests that, while a phytoene boost might start the initial steps of the chromoplast differentiation program, an enhanced production of downstream carotenoids is essential to complete it.

**Figure 5.**
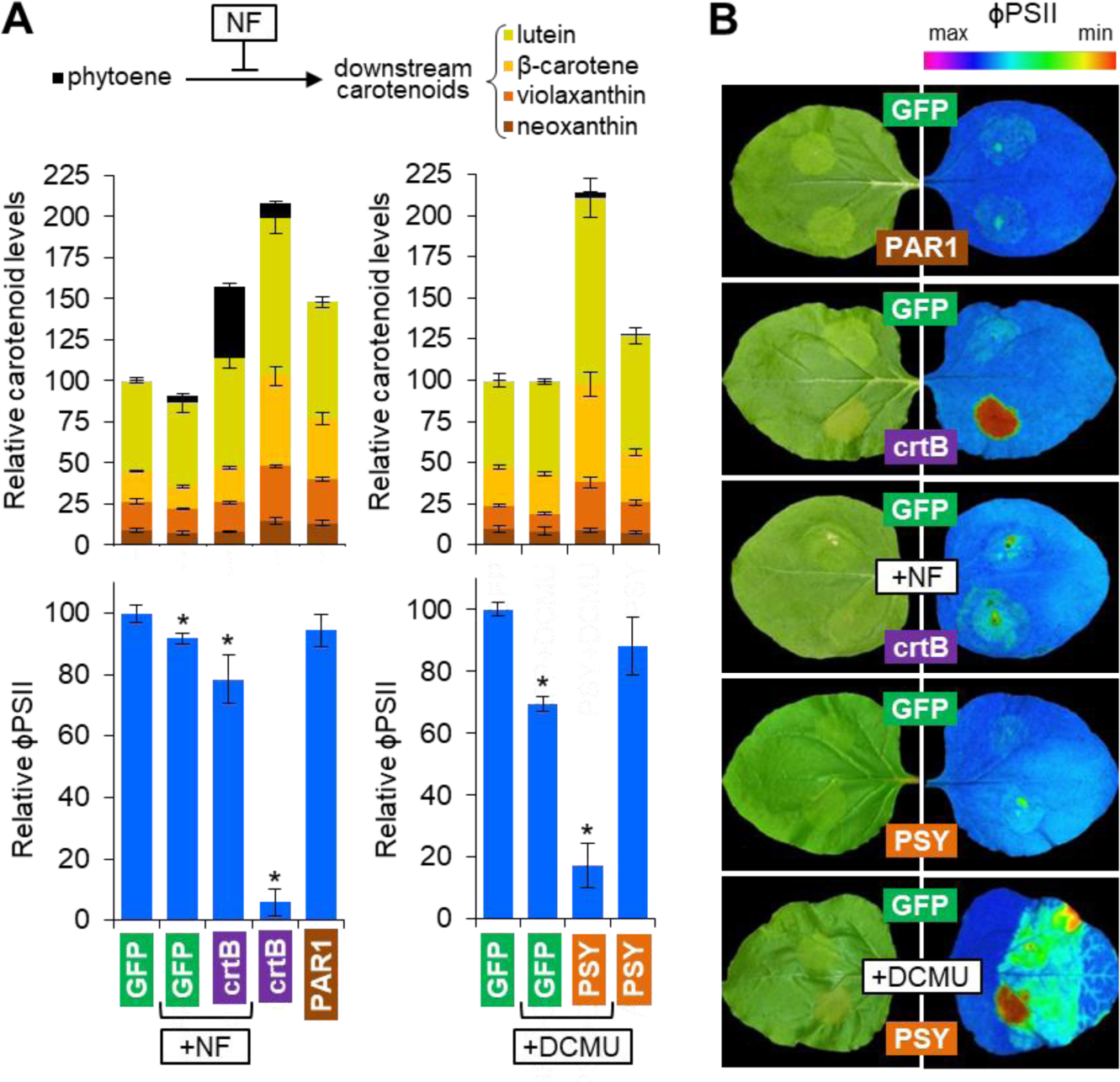
Transformation of leaf chloroplasts into chromoplasts requires a reduction of photosynthetic capacity and production of carotenoids downstream of phytoene. A, Carotenoid levels and ϕPSII in leaves 96 hours after agroinfiltration with the indicated constructs. In all cases, plot values correspond to mean and standard deviation values of n=3 independent samples. Asterisks in ϕPSII plots mark statistically significant changes relative to untreated GFP controls (*t* test, *P* < 0.05). Samples infiltrated with NF at 24 hpi or treated with DCMU 24 h before agroinfiltration are indicated. B, representative images of agroinfiltrated leaves at 96 hpi and their corresponding chlorophyll fluorescence for ϕPSII.

### Pharmacological inhibition of photosynthetic activity reduces the phytoene threshold to initiate chloroplast-to-chromoplast transition in leaves

In contrast with the results using crtB, overexpression of PSY-encoding genes from *Arabidopsis* and tomato could not elicit the characteristic yellow leaf phenotype associated with chromoplast differentiation (Fig. 5 and Fig. S7) (Fraser et al., 2007; Maass et al., 2009). Interestingly, the plant enzymes yielded significantly lower levels of phytoene compared to crtB and did not substantially impact photosynthesis as deduced from ϕPSII values (Fig. 5 and Fig. S7). We hence speculated that phytoene might act as a metabolic threshold switch that only alters photosynthetic performance of chloroplasts when exceeding a certain level. To overcome this putative threshold, we treated *N. benthamiana* leaves with DCMU (3-(3,4-dichlorophenyl)-1,1-dimethylurea, diuron), a widely used inhibitor of photosynthesis that interrupts the photosynthetic electron transport chain. The next day, treated and untreated control leaves were agroinfiltrated with constructs encoding either GFP or the *Arabidopsis* PSY enzyme. At 96 hpi, DCMU-treated GFP leaf sections showed a decrease in ϕPSII but unchanged carotenoid levels compared to untreated samples (Fig. 5). By contrast, DCMU-treated PSY sections showed a more dramatic drop in ϕPSII and much higher levels of carotenoids than untreated PSY or GFP controls (Fig. 5A). As a consequence, PSY leaf sections treated with DCMU turned yellow (Fig. 5B), similar to that observed when chloroplast-to-chromoplast differentiation was triggered by crtB in the absence of inhibitor. In summary, our results are consistent with the existence of a two-step process responsible for the crtB-mediated transformation of leaf chloroplasts into chromoplasts (Fig. 6). Firstly, chloroplast identity is weakened by overaccumulation of phytoene and, secondly, increased production of carotenoids in pre-conditioned chloroplasts allows the differentiation of chromoplasts. Without pre-conditioning, carotenoid levels can increase but chromoplasts do not differentiate, as shown in untreated leaves producing PAR1 or PSY. Moreover, if carotenoids downstream of phytoene are not produced, pre-conditioned chloroplasts remain unchanged, as shown in NF-treated (p)crtB leaves (Fig. 5).

**Figure 6.**
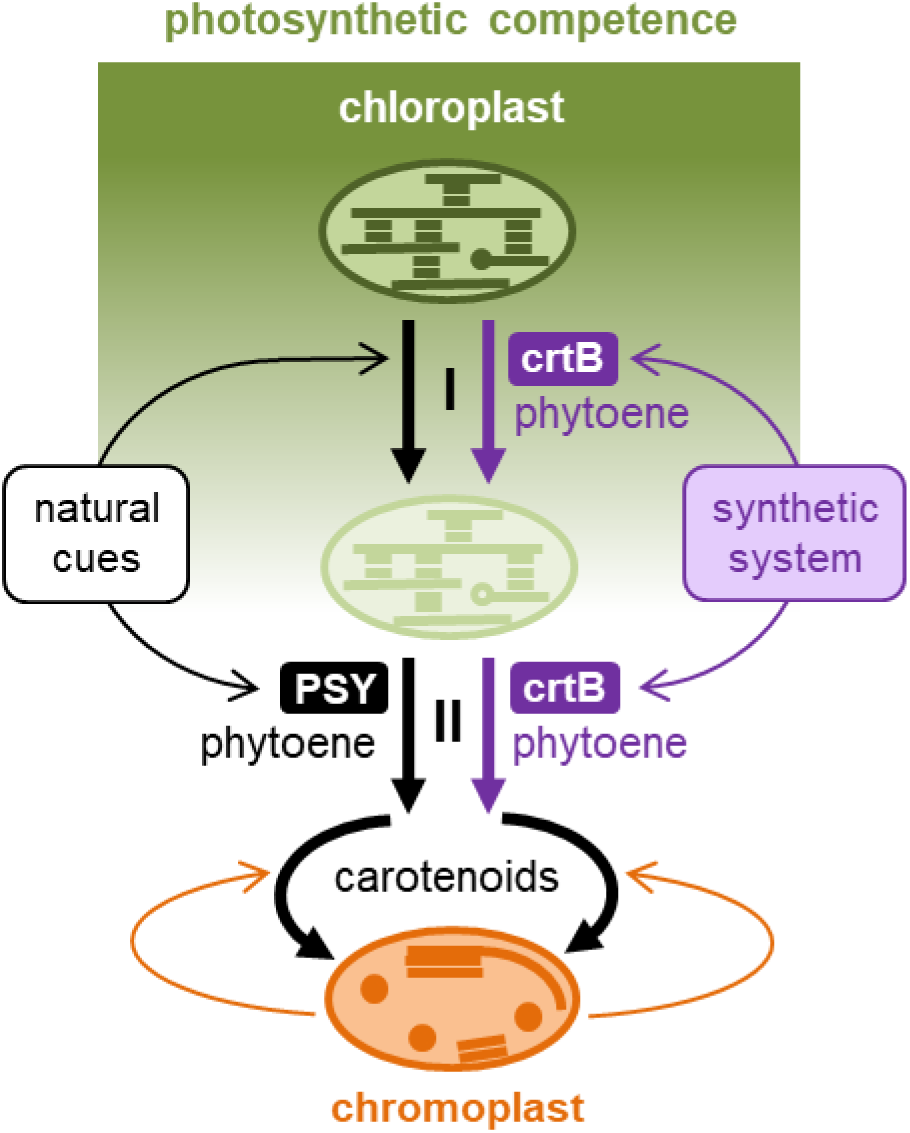
Model of the chloroplast-to-chromoplast differentiation process. Plant developmental programs create organs with different degrees of photosynthetic capacity and hence chloroplast identity, from strong (e.g., leaves) to weak (e.g., green fruits) or absent (e.g., roots). Loss of chloroplast identity appears to be the first phase (I) in chromoplast differentiation. In a second phase (II), developmental cues promote the expression of genes encoding PSY and other carotenoid biosynthetic enzymes. Enhanced production of carotenoids then reprograms plastid-to-nucleus communication, changes plastidial ultrastructure, and results in the differentiation of chromoplasts, which in turn promotes biosynthesis and improves storage of carotenoids. The two phases can be synthetically engineered in leaves by overproducing phytoene using crtB. When phytoene exceeds a certain level, it interferes with the photosynthetic capacity of leaf chloroplasts. This acts as a metabolic switch that allows the formation of chromoplasts after phytoene is converted into downstream carotenoids by endogenous enzymes.

## Discussion

The regulation of plastid identity is a core process in plants that remains poorly defined. Our work shows that chromoplasts can be synthetically differentiated from leaf chloroplasts in all plants tested, despite this is something that only a few species can do in Nature. This synthetic system has allowed us to propose a model for chromoplast differentiation that also applies to natural systems (Fig. 6). In a first phase (I), chloroplasts must become competent (i.e., pre-conditioned) by lowering their photosynthetic capacity, whereas increased production of carotenoids completes the differentiation of chromoplasts in a second phase (II).

In our synthetic system, phase I was very fast (hours) and required a sufficient amount of phytoene to break chloroplast identity in leaves. In chloroplasts, carotenoids such as lutein, beta-carotene, violaxanthin, and neoxanthin are required to maintain the properties of photosynthetic membranes and pigment– protein complexes responsible for harvesting sunlight and transferring excitation energy to the photosystems (Domonkos et al., 2013; Liguori et al., 2017; Rodriguez-Concepcion et al., 2018). Phytoene is not normally detected in leaf chloroplasts as it is readily converted into downstream (photosynthesis-related) carotenoids. Overaccumulation of phytoene (or a phytoene derivative) might somehow compete with endogenous carotenoids for their binding to photosynthetic protein complexes and membranes, interfere with their functions, and eventually cause the changes that we observed in photosynthetic competence. Consistent with this hypothesis, engineered accumulation of non-chloroplast carotenoids such as astaxanthin in plants alters the properties of thylakoids and grana and interferes with the photosynthetic machinery at several other levels (Roding et al., 2015; Liguori et al., 2017). Transplastomic tobacco plants producing massive levels of asthaxanthin developed leaf plastids that lost their chloroplast features (Lu et al., 2017), similar to our results

In nature, chloroplasts might become competent for differentiation into chromoplasts without the need of a phytoene boost. In tomato, the chloroplasts of green fruit are much less photosynthetically active than those of leaves (Cocaliadis et al., 2014; Lado et al., 2016). Thus, tomato green fruit and other organs, tissues or/and developmental stages in which chloroplast identity is weak or non-existent (e.g., in dark-grown calli, tubers, or roots) might be considered as “naturally competent” to differentiate chromoplasts when carotenoid biosynthesis is upregulated. In agreement, overproduction of plant PSY enzymes resulted in chromoplast-like structures arising in green tomato fruit and non-photosynthetic *Arabidopsis* tissues but had no effect on leaves (Fraser et al., 2007; Maass et al., 2009) unless previously conditioned by shutting down their photosynthetic capacity (Fig. 5). Similarly, upregulation of OR triggers carotenoid overaccumulation and chromoplast differentiation in tomato fruit, potato tubers, cauliflower curds, or *Arabidopsis* calli, but not in the leaves of any of these plants (Li et al. 2001; Lopez et al. 2008; Yuan et al. 2015; Yazdani et al. 2019). Interestingly, OR does not appear to be required for crtB-mediated differentiation of leaf chloroplasts into chromoplasts (Fig. S2). OR has been shown to promote PSY activity and stability, and some OR mutants prevent carotenoid (particularly β-carotene) metabolism (Tzuri et al., 2015; Zhou et al., 2015; Sun et al., 2018; Welsch et al., 2018). Our results suggest that no extra function would be required for OR upregulation to cause chromoplast development from plastids with weak or no chloroplast identity.

In phase II, which likely overlaps with phase I, carotenoid accumulation occurs concomitantly with the remodeling of the internal plastid structures, with both factors synergistically activating each other (Fig. 6). Strikingly, in agroinfiltrated *N. benthamiana* leaves this process takes place without changes in the amount of chlorophylls (Fig. 2), which might remain trapped in the membranous structures observed in the chromoplasts (Fig. 1). These results confirm that chlorophyll breakdown and chromoplast differentiation are independent processes, as already shown in mutants such as tomato *green flesh* (*gf*), in which impairment of chlorophyll degradation during fruit ripening has no effect on the formation of chromoplast membranes and the accumulation of carotenoids (Cheung et al., 1993). The new structures created following the disassembly of photosynthetic grana and thylakoids (Fig. 1) likely contribute to reaching high carotenoid levels by accommodating increasing amounts of carotenoids and by preventing their degradation (Egea et al., 2010; Jarvis and Lopez-Juez, 2013; Lado et al., 2016; Llorente et al., 2017; Sun et al., 2018; Sadali et al., 2019). They might additionally enhance carotenoid production by stimulating the activity of endogenous carotenoid biosynthetic enzymes (including PSY), many of which are membrane-associated (Ruiz-Sola and Rodriguez-Concepcion, 2012). Indeed, NF treatment reduced the total amount of carotenoids that accumulated in (p)crtB samples (Fig. 5A), suggesting that chromoplast differentiation was linked to enhanced metabolic flux into and through the carotenoid pathway. The observation that crtB-producing leaves virtually double their carotenoid contents compared to GFP controls (Fig. 2, 4 and 5) with only minor changes in the expression of biosynthetic genes (Fig. S4) further supports the conclusion that post-translational effects on enzyme activity or/and improved carotenoid storage and stability are responsible for the carotenoid overaccumulation phenotype.

The structural changes associated with the chloroplast-to-chromoplast transformation involve reorganization of the plastidial proteome (Fig. 1E) but also a global reprogramming of nuclear gene expression (Fig. 3) and primary metabolism (Fig. S5). It is likely that implementing these changes relies on retrograde signals produced by differentiating plastids. Carotenoid degradation can generate signaling molecules that regulate many developmental processes in plants, including plastid development (Hou et al., 2016; Wang et al., 2019). The observation that *Arabidopsis ccd1 ccd4* mutants defective in carotenoid cleavage dioxygenase activity in the cytosol (via CCD1) and the plastids (via CCD4) were not affected in the crtB-dependent leaf phenotype (Fig. S2) suggests that signals independent of CCDs or carotenoids are responsible for eliciting the changes in nuclear gene expression and cell metabolism supporting chromoplast biogenesis. The specific nature of such signals remains to be discovered.

In summary, we show that chromoplast differentiation only requires metabolic cues (i.e., enough phytoene and downstream carotenoid production). While our conclusions are based on a synthetic system (i.e., the expression of a bacterial gene in leaf cells), the similarity of the transcriptomic profiles between this process and fruit ripening (Fig. 3) strongly supports that this is a basic general mechanism for chloroplasts to become chromoplasts. In nature, however, developmental cues play a fundamental role by making chloroplasts competent (phase I) and by regulating the expression of carotenoid biosynthetic genes (phase II). Signals produced by differentiating plastids are also hardwired to the process as they support the organellar transformation by reprogramming nuclear gene expression and whole cell metabolism. Besides serving to successfully address a long-standing question in plant biology (i.e., plastid identity), the very simple and straightforward tool that we describe here to induce leaf chloroplast-to-chromoplast differentiation on demand should allow to improve the nutritional quality of leaf crops by creating an organellar sink for the accumulation of carotenoids and other plastidial phytonutrients (Lado et al., 2016; Giuliano, 2017; Llorente et al., 2017; Sun et al., 2018; Wurtzel, 2019).

## Materials and methods

### Plant material and growth conditions

*Nicotiana tabacum* Xanthi nc, *Nicotiana benthamiana* RDR6i, and *Arabidopsis thaliana* Columbia-0 (Col) and Landsberg *erecta* (Ler) plants were grown under standard conditions as described previously (Llorente et al., 2016; Majer et al., 2017). Growth of double mutants *ccd1 ccd4* (Zhou et al., 2015) and *ator atorl* (Schaub et al., 2018), both in the Col background, was facilitated by transferring them to low light (40 μmol photons m^−2^ s^−1^) days after germination. For generating dark-induced leaf senescence, detached leaves were maintained inside dark, humid chambers until visible yellowing occurred. Collected plant material was frozen in liquid nitrogen, lyophilized, and then homogenized to a fine powder using a TissueLyser system (Qiagen) for further analyses.

### Transmission electron microscopy

Transmission electron microscopy of plant leaves was performed as previously described (D’Andrea et al., 2018).

### Gene constructs

Viral vectors used in this study have been described (Majer et al., 2017) with the following exceptions. A plasmid with recombinant clone TEV-GFP was constructed from plasmid pGTEVa (Bedoya et al., 2012) and used for *Arabidopsis* Ler plants. The binary viral vector pGTuMV-UK1 was used for *Arabidopsis* Col plants. Briefly, a cDNA of the UK1 isolate of *Turnip mosaic virus* (TuMV; GenBank accession number NC_002509.2) was cloned in vector pG35Z (Cordero et al., 2017) flanked by the *Cauliflower mosaic virus 35S* promoter and terminator.

To generate the different crtB versions, we amplified by PCR the *Pantoea ananatis crtB* gene (Majer et al., 2017). We also amplified cDNA sequences encoding the plastid-targeting peptide of *Arabidopsis* hydroxymethylbutenyl 4-diphosphate synthase (Gas et al., 2009), *Arabidopsis* PSY and tomato PSY2. Primers used for cloning procedures are detailed in Table S6. PCR products were cloned using the Gateway system first into plasmid pDONR-207 and then into plasmid pGWB405 (Nakagawa et al., 2007) to generate 35S:crtB-pGWB405, 35S:(p)crtB-pGWB405, 35S:(p)crtB-GFP-pGWB405, 35S:AtPSY-pGWB405, and 35S:SlPSY2-pGWB405, or plasmid pGWB506 (Nakagawa et al., 2007) to generate 35S:GFP-crtB-pGWB506. To generate TuMV-crtB, the crtB sequence was cloned by Gibson assembly between positions 8758 and 8759 of pGTuMV-UK1 after amplification with flanking sequences complementing the TuMV nuclear inclusion a protease (NIaPro) cleavage site (Table S6) to mediate the release of the crtB protein from the viral polyprotein.

### Transient expression assays

For transient expression studies using viral vectors, leaves of 4 to 6 week-old *N. tabacum* and *Arabidopsis* plants were mechanically inoculated with crude extracts from frozen-stored infected plant tissue and collected upon the appearance of the yellowing phenotype as described previously (Majer et al., 2017). For agroinfiltration experiments, the second or third youngest leaves of 4 to 6 week-old *N. benthamiana* plants were infiltrated with *A. tumefaciens* strain GV3101 carrying plasmids of interest following the procedure described previously (Sparkes et al., 2006). Gene silencing was prevented by co-agroinfiltration with a strain carrying the helper component protease (HcPro) of the watermelon mosaic virus (WMV) in plasmid HcProWMV-pGWB702 (kindly provided by Juan José López-Moya and Maria Luisa Domingo-Calap). Infiltration cultures were grown on LB medium at 28°C and used at optical density at 600 nm (OD600) of 0.5. For pharmacological treatments, norflurazon (NF) or diuron (DCMU) were diluted in water and 0.05 % Tween 20. The treatments with NFZ (2 µM) were performed by infiltration with a syringe of leaf areas that had been agroinfiltrated with different constructs 24 h earlier. DCMU (10 µM) was applied on the leaf surface with a fine paintbrush 24 h before agroinfiltration.

### Transcript analyses

Total RNA was extracted from leaves with the Maxwell 16 LEV Plant RNA Kit (Promega) and quantified with a NanoDrop (Thermo Scientific) as described (Majer et al., 2017). For reverse transcription-quantitative PCR (RT-qPCR) analyses, the First Strand cDNA Synthesis Kit (Roche) was used to generate cDNA according to the manufacturer’s instructions, with anchored oligo(dT)_18_ primers and 1 μg of total RNA. Relative mRNA abundance was evaluated via quantitative PCR using LightCycler 480 SYBR Green I Master Mix (Roche) on a LightCycler 480 real-time PCR system (Roche). Gene expression analysis of *N. tabacum SAG12* gene was conducted using primers NtSAG12_qPCR-F (5’-A T T C A T G G G G C A G T A A A T G G-3’) and NtSAG12_qPCR-R (5’-G A A G C G T C C A T A G C A A G T C C-3’) and the L25 ribosomal protein gene for normalization (Schmidt and Delaney, 2010) using primers NtRPL25_qPCR-F (5’-C A A G G C A C A G G C A G C T A A G G-3’) and NtRPL25_qPCR-R (5’-A G G T C G G T G G A A T G T A A C T T T T G-3’).

RNAseq service was performed by Sequentia Biotech SL (Barcelona, Spain). RNA concentration in each sample was assayed with a ND-1000 spectrophotometer (NanoDrop) and its quality assessed with the TapeStation 4200 (Agilent Technologies). Indexed libraries were prepared from 1 μg/ea purified RNA with TruSeq Stranded mRNA Sample Prep Kit (Illumina) according to the manufacturer’s instructions. Libraries were quantified using the TapeStation 4200 and pooled such that each index-tagged sample was present in equimolar amounts, with final concentration of the pooled samples of 2 nM. The pooled samples were subject to cluster generation and sequencing using a NextSeq 500 System (Illumina) in a 2×75 paired end format at a final concentration of 1.8 pmol. The raw sequence files generated (.fastq files) underwent quality control analysis using FastQC (http://www.bioinformatics.babraham.ac.uk/projects/fastqc/). Data analysis was performed with the online platform AIR (www.transcriptomics.cloud) using the SolGenomics Network (https://solgenomics.net/) *N. benthamiana* 1.01 (Niben v101) reference genome.

### Protein extraction and immunoblot analyses

Protein extraction, quantification, and immunoblot analyses were performed as described (Pulido et al., 2016) using anti-fibrillin (Simkin et al., 2007) or anti-PsbA serum (Agrisera).

### Metabolite analyses

Carotenoids were analyzed as previously described (Llorente et al., 2016). Phytoene was quantified using a concentration curve with a commercial standard (Sigma). Primary metabolites were extracted as described previously (Lisec et al., 2006) using approximately 20 mg of lyophilized leaf tissue. Derivatization and gas chromatography-time of flight-mass spectrometry (GC-TOF-MS) analyses were carried out as described previously (Lisec et al., 2006). Metabolites were identified manually following use of TagFinder software in combination with the reference library mass spectra and retention indices from the Golm Metabolome Database, http://gmd.mpimp-golm.mpg.de. The parameters used for the peak annotation of the 52 metabolites can be found in Table S4, which follows recommended standards on the report of metabolite data (Fernie et al., 2011). Data were normalized to the mean value of the GFP control samples (i.e., the value of all metabolites for GFP control samples was set to 1). The means and standard errors of five to six replicates at 96 hpi are presented in Table S5.

### Photosynthetic measurements

Photosynthetic efficiencies were assessed by measuring chlorophyll *a* fluorescence with a MAXI-PAM fluorometer (Heinz Walz GmbH). Leaves were placed under the camera and effective quantum yield (ΔF/Fm’) was measured as (Fm’−Fs)/Fm’, where Fm’ and Fs are, respectively, the maximum and the minimum fluorescence of light exposed plants. The light intensity chosen was 21 PAR (actinic light, AL=2) as the last value able to generate a response in the (p)crtB-infiltrated areas before having a null photosynthetic activity. Each value is the average result of three biological replicates and three different AOI for each replicate. NPQ was also measured using the MAXI-PAM unit. All recordings were performed every day at the same time slot, but the order of the samples was randomized in order to reduce the bias related to the length of the light stress recovery protocol. Plants were dark-adapted for 30 min before measurement and then submitted to a continuous 801 PAR light (AL=17) for 10 min. After this period plants were left recovering in darkness for 40 min. During the whole protocol, Ft was monitored and Fm’ values were estimated with a saturating pulse (SAT) every 60 seconds. NPQ and its relative components qE, qZ and qI were calculated as described (Coate et al., 2013) with some modifications. Briefly: NPQ was calculated as (Fm-Fm_0_)/Fm, where Fm and Fm_0_ are the maximum fluorescence after the dark acclimation and after the light stress, respectively. The relative contributions of qE, qZ and qI to NPQ were estimated by monitoring NPQ relaxation kinetics in the dark: following the 10 min exposure to saturating light used to measure NPQ, leaves were left in darkness, and Fm_0_ was measured again after 10 and 40 min. The qE component of NPQ relaxes within 10 min of a leaf being placed in darkness such that NPQ persisting after 10 min in the dark consists of qZ + qI. The qZ component of NPQ relaxes within tens of minutes so that NPQ persisting after 40 min in the dark (when the Ft value is linear) consists of qI, which is either irreversible in the dark or requires several hours to relax. Consequently, (qI+qZ) was calculated as (Fm-Fm_1_)/Fm_1_, where Fm1 is the value of Fm measured after 10 min in the dark following NPQ measurement. qZ was calculated as (Fm-Fm2)/Fm2, where Fm2 is the value of Fm measured after 40 min in the dark following measurement of NPQmax. qE was calculated as NPQ–(qI+qZ) and qI was calculated as (qI+qZ)-qZ. For the calculation of the de-epoxidation state (DES), agroinfiltrated leaf areas were exposed for 10 min to a continuous 801 PAR light (AL=17) in the MAXI-PAM unit, sampled under the same light and immediately frozen before pigment extraction and quantification. The operation of the xanthophyll cycle, comprising the sequential de-epoxidation of the pigments violaxanthin (Vx) to antheraxanthin (Ax) and zeaxanthin (Zx) was followed by calculating DES as (Zx+0.5×Ax)/(Zx+Ax+Vx), where Zx, Ax and Vx are the concentrations of the corresponding xanthophylls.

### Respiration measurements

Before respiration measurements, *N. benthamiana* plants were placed in the dark for about 30 min to avoid light-enhanced dark respiration. Four leaf discs of 3.8 cm^2^ each were harvested from leaf sections of two different plants infiltrated with the (p)crtB construct, weighted and placed into the respiration cuvette containing the respiration buffer (30 mM MES pH 6.2, 0.2 mM CaCl_2_). Oxygen uptake rates were measured in darkness using a liquid-phase Clark-type oxygen electrode (Rank Brothers) at a constant temperature of 25ºC. Dry weights from leaf discs were determined after drying for 2 days at 60ºC.

### Statistical analyses

Differentially expressed genes (DEGs) were identified by comparing crtB and GFP RNAseq datasets with the DESeq2 statistical method in the AIR platform. The resulting crtB/GFP list was filtered using cut-offs of FDR <0.05 and log2-transformed fold change, log2FC, >0.585 for upregulated genes and <-0.599 for downregulated genes. Gene ontology enrichments were identified by the Parametric Analysis of Gene Set Enrichment (PAGE) function of the AgriGO v2.0 web-based tool (http://bioinfo.cau.edu.cn/agriGO/) after transforming the gene IDs to the Niben v04.4 annotation. For the comparison of different biological systems, we selected the significantly enriched gene ontologies from our *N. benthamiana* RNAseq experiment (p and q values <0.05) and compared their Z-score values with those obtained from the analysis of published RNAseq data of tomato fruit ripening and *Arabidopsis* leaf senescence. In particular, we used the RPM values of the total pericarp at mature green (MG), light ripe (LR) and red ripe (RR) stages (Shinozaki et al., 2018) and the FPKM values of the 16D and 30D senescence stages (Woo et al., 2016). DEGs resulting from LR/MG, RR/MG and 30D/16D comparisons were filtered as described above for *N. benthamiana* crtB/GFP.Student’s *t*-tests were used for the rest of statistical analyses using GraphPad Prism 5.0a (GraphPad Software).

## Acknowledgements

We greatly thank Jaume F. Martinez-Garcia for fruitful discussions on the manuscript, Ralf Welsch and Li Li for providing seeds of the *Arabidopsis ccd1 ccd4* and *ator atorl* mutants, respectively, Juan José López-Moya and Maria Luisa Domingo-Calap for the gift of the HcProWMV-pGWB702 plasmid, and M. Rosa Rodriguez-Goberna for excellent technical support. The help of Martí Bernardo, Fidel Lozano, Lídia Jiménez and members of the CRAG core facilities is also appreciated. This work was funded by the European Regional Development Fund (FEDER) and the Spanish Agencia Estatal de Investigación (grants BIO2017-84041-P, BIO2017-83184-R, BIO2017-90877-REDT, and BES-2017-080652), Ministry of Education, Culture and Sports (AP2012-3751 and FPU16/04054), and Generalitat de Catalunya (2017SGR-710). We also thank the financial support of the European Union’s Horizon 2020 (EU-H2020) COST Action CA15136 (EuroCaroten) and Marie S. Curie Action 753301 (Arcatom), the Severo Ochoa Programme for Centres of Excellence in R&D 2016-2019 (SEV-2015-0533) and the Generalitat de Catalunya CERCA Programme to CRAG. BL is supported by grants from the CSIRO Synthetic Biology Future Science Platform and Macquarie University. LM is supported by La Caixa Foundation PhD INPhINIT fellowship LCF/BQ/IN18/11660004, which received funding from the EU-H2020 (MSCA grant 713673). ARF is supported by the Deutsche Forschungsgemeinschaft (DFG TRR 175)

