## SUPPLEMENTAL FIGURES for "Synthetic biogenesis of chromoplasts from leaf chloroplasts"

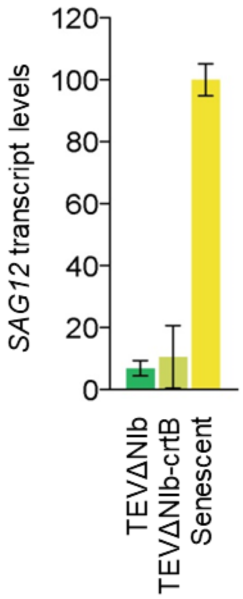


**Figure S1. *SAG12* gene expression in *N. tabacum* leaves.** Transcript levels were quantified by RT-qPCR in symptomatic leaves from plants carrying the indicated viral vectors at 7 dpi. As a control, *SAG12* expression was also tested in non-inoculated leaves that were harvested from the plant and kept in the dark for 10 days (senescent).

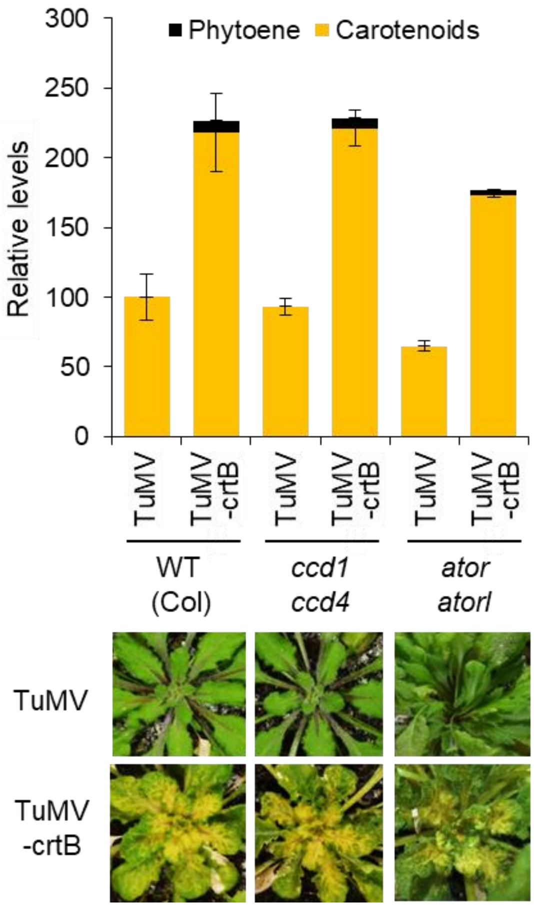


**Figure S2. Carotenoid cleavage and OR activity are not required for the crtB-associated phenotype.** Arabidopsis Col wild-type (WT) and double mutant plants were grown under short day conditions (8h of low light and 16h of darkness) for 2 weeks (WT and *ccd1 ccd4*) or 5 weeks (*ator atorl*) and then inoculated with the indicated viral vectors. Pictures and samples for carotenoid analysis were taken 17 days after inoculation. Plot shows the mean and standard deviation of n=3 independent samples. Carotenoid levels are represented relative to those in WT samples inoculated with the empty vector control (TuMV).

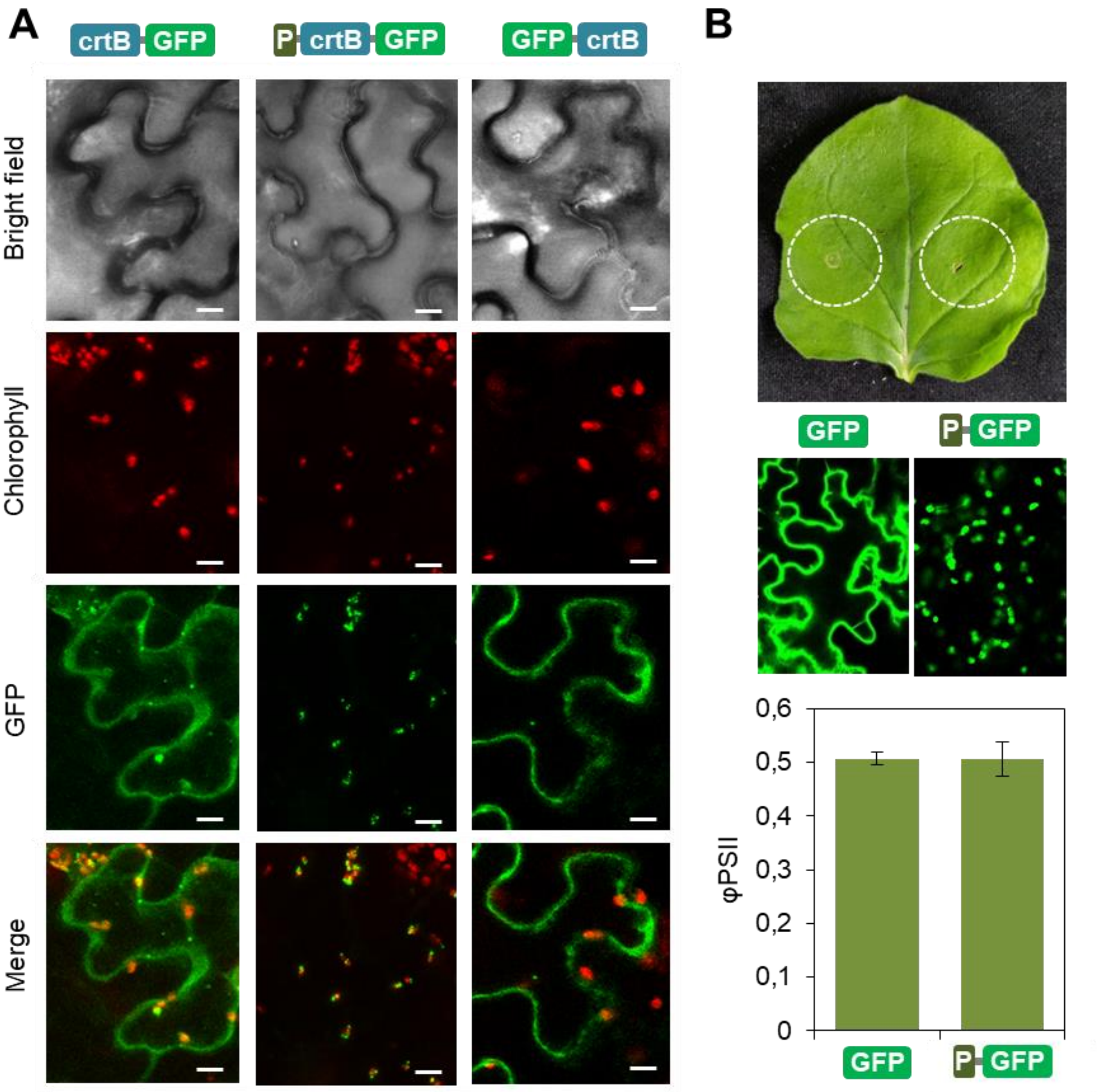


**Figure S3. Localization of GFP-tagged proteins.** A, Subcellular localization of the indicated constructs in agroinfiltrated *N. benthamiana* leaf cells by confocal laser scanning microscopy. Pictures were taken 3 days after agroinfiltration (72 hpi). Bars, 20 µm. B, *N. benthamiana* leaf agroinfiltrated with constructs to express a plastid-targeted version of GFP (P-GFP) or the untargeted GFP protein (upper picture). At 72 hpi, GFP fluorescence was examined in agroinfiltrated tissue to confirm the localization of P-GFP in chloroplasts and GFP in the cytosol (middle pictures). Effective quantum yield of PSII was quantified at 96 hpi (lower plot).

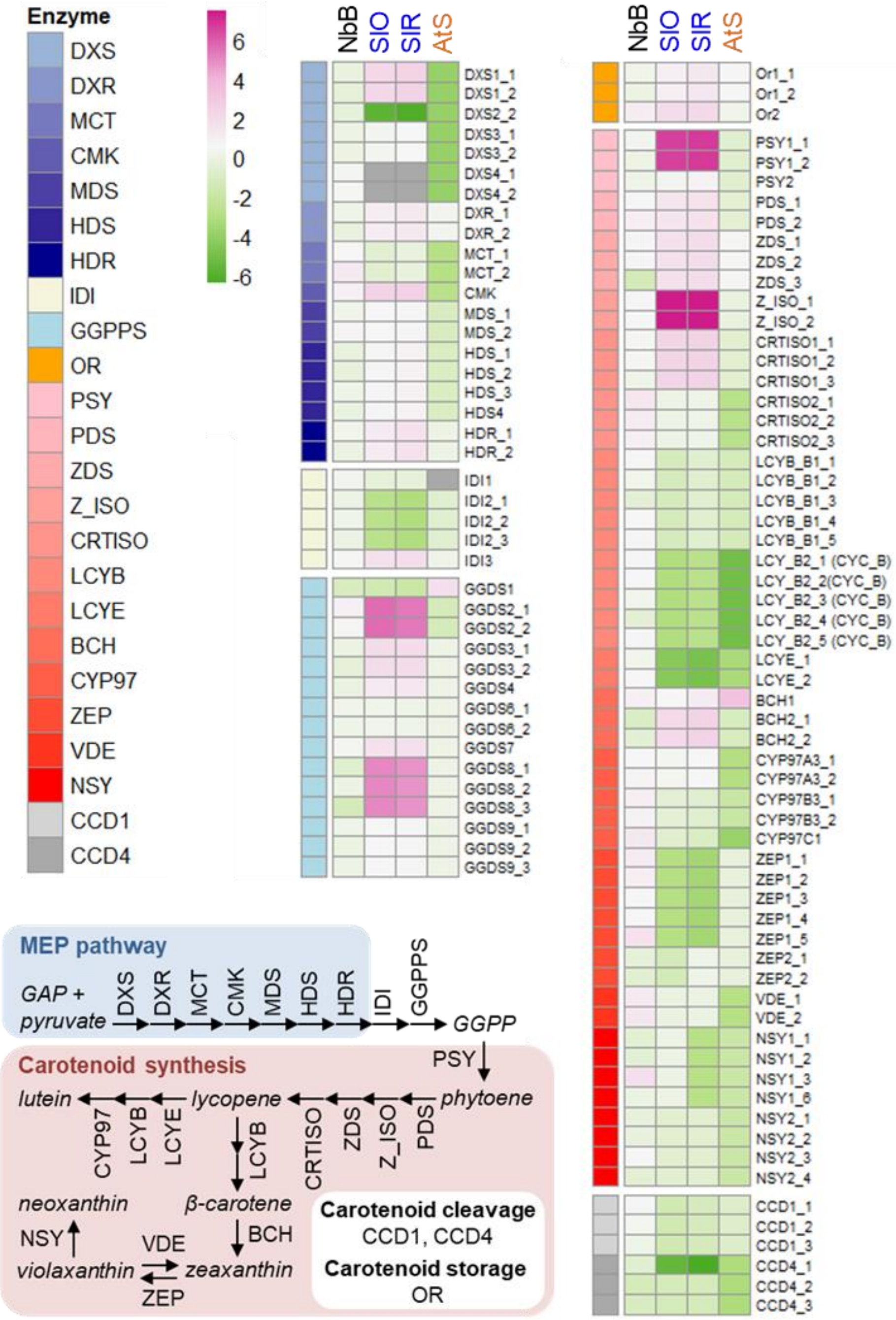


**Figure S4. Level of transcripts for carotenoid-related proteins.** Heatmap represents log2FC values in *N. benthamiana* leaves 4 days after agroinfiltration with (p)crtB compared to GFP (lane NbB). Data publicly available from tomato fruit (light ripe vs. mature green, SlO, and red ripe vs. mature green, SlR) and Arabidopsis leaves (senescent -30D- vs. controls -16D-, AtS) are also shown. The position of enzymes in pathways and/or the biological function of the proteins selected are indicated in the cartoon and represented by colors.

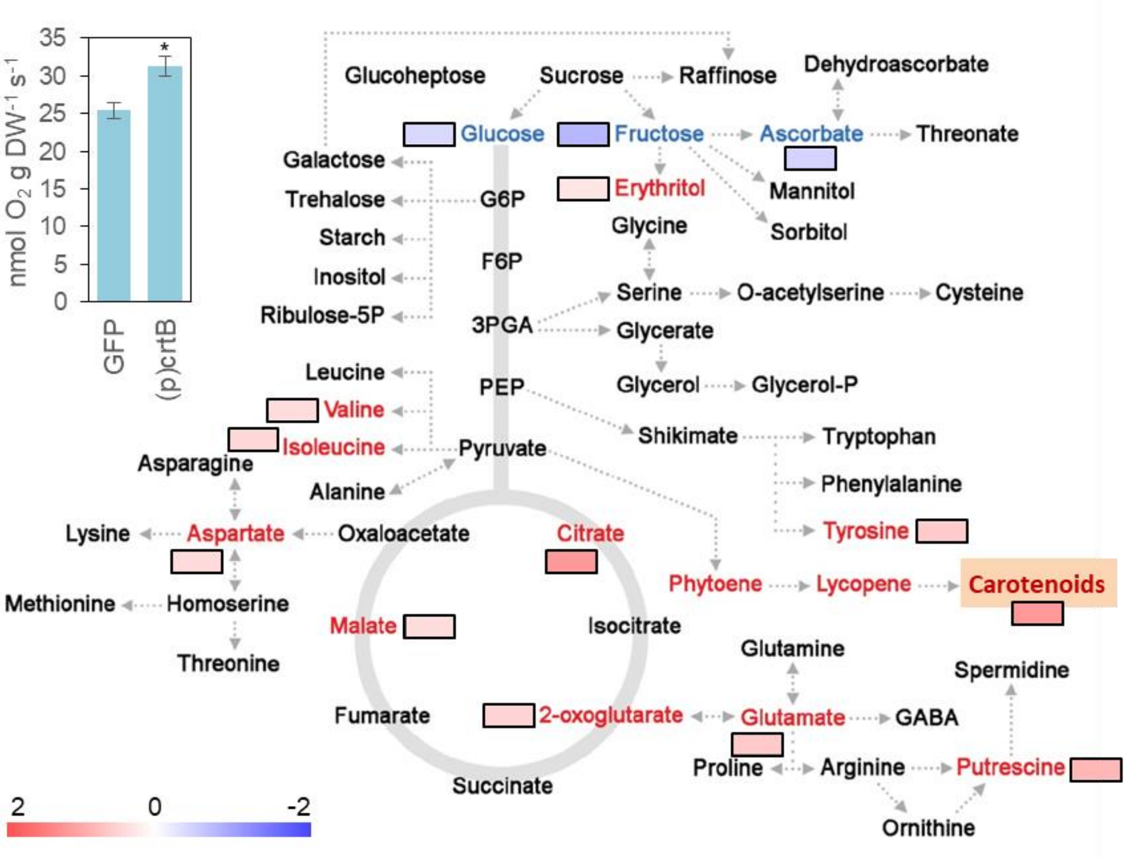


**Figure S5. Chromoplast differentiation induces glycolytic and oxidative energy metabolism.** Heatmap represents statistically significant FC (fold-change) values of metabolite levels in *N. benthamiana* leaves 4 days after agroinfiltration with (p)crtB relative to those in GFP controls. Inset represents respiration rates. Mean and standard deviation values of n = 3 independent samples are shown. Asterisk represents statistical significance (*t* test, $P < 0.05$).

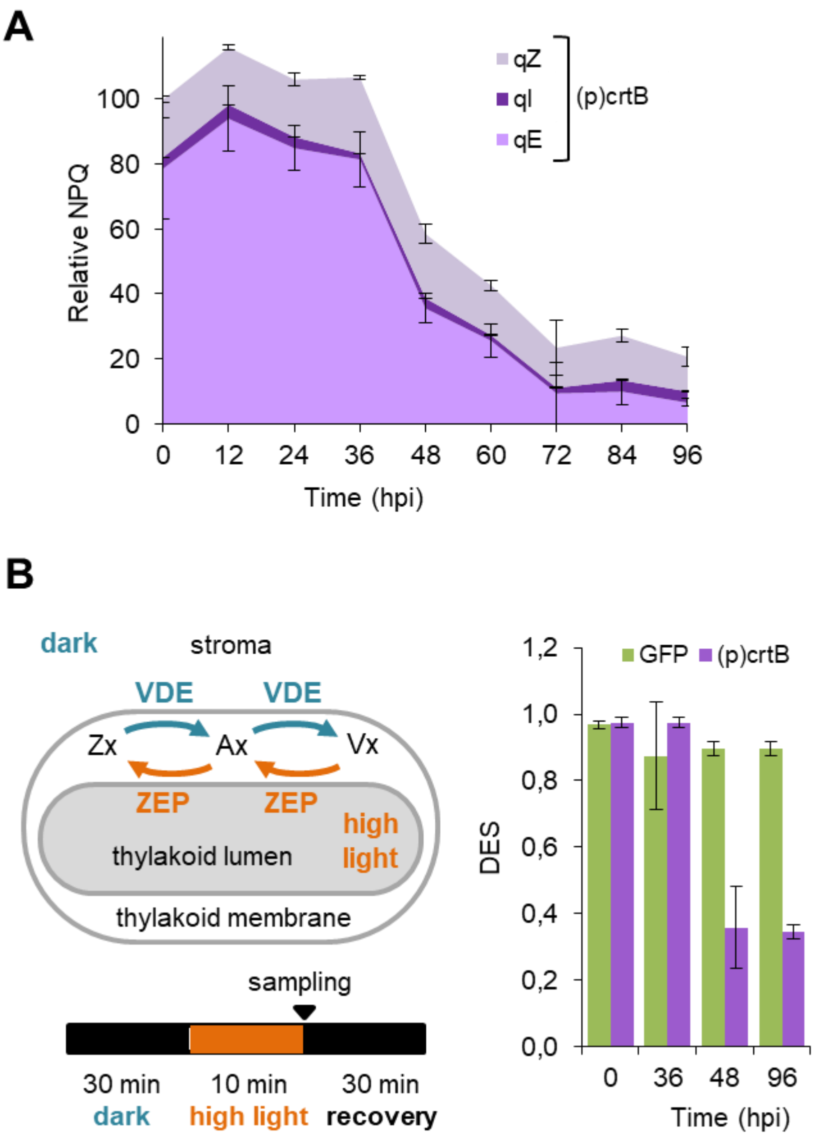


**Figure S6. Chromoplast differentiation in leaves reduced NPQ and impairs the xanthophyll cycle.** A, Non-photochemical quenching (NPQ) components in *N. benthamiana* leaf sections agroinfiltrated with (p)crtB. B, Leaves producing GFP or (p)crtB were treated as shown in the left panel and then collected to quantify their carotenoid levels. De-epoxidation state (DES) was calculated as (Zx+0.5×Ax)/(Zx+Ax+Vx), where Zx, Ax and Vx are the concentrations of zeaxanthin, antheraxanthin, and violaxanthin, respectively.

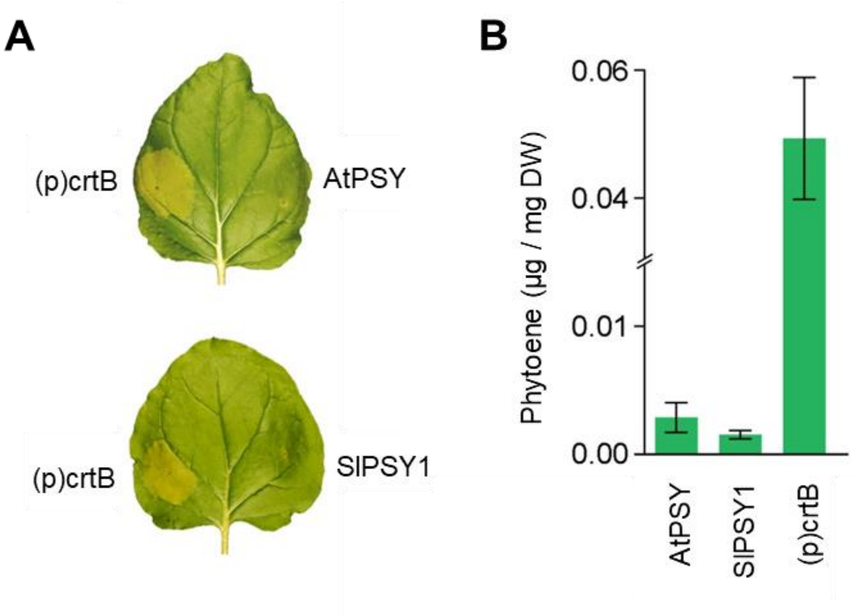


**Figure S7. Plant PSY enzymes do not induce chromoplast differentiation in leaves.** A, Phenotypes of *N. benthamiana* leaf sections producing the indicated proteins at 96 hpi. B, Phytoene content in the leaf sections shown in A.
